# Theoretical Analysis of Power-Law Stress Relaxation and Calcium-Dependent Passive Mechanics in Cardiac Muscle

**DOI:** 10.1101/2025.02.26.640338

**Authors:** Filip Ježek, Anthony J. Baker, David A. Nordsletten, Daniel A. Beard

## Abstract

This study investigates the passive viscoelastic mechanical properties of cardiac muscle by introducing a theoretical model that explains the power-law kinetics of passive stress decay. The model accounts for two parallel processes contributing to passive mechanics: an elastic component and a viscoelastic component designed to simulate stress/strain-mediated unfolding of serial domains in the titin molecule. Under stress, serial globular domains within the elastic region of the titin molecule reversibly unfold. This unfolding phenomenon contributes to both hysteresis (a lag in stress between loading and unloading) and preconditioning effects in simulated striated muscle mechanics. Moreover, experimental evidence indicates that stress relaxation in cardiac muscle follows a power law, and that the muscle’s nonlinear stress-strain relationship and hysteresis behavior are calcium-dependent. To analyze these mechanical phenomena, we simulate the apparent viscous element as a mesoscopic-scale ensemble of chains, each composed of serial globular domains that unfold in a stress-dependent manner. Although the model was developed to represent the behavior of titin, it equivalently represents any contributing process involving a linked series of domains that undergo stress-mediated unfolding. By providing a unified basis for the observed viscoelastic and preconditioning effects, calcium dependency, and power-law stress relaxation phenomena, this study offers a novel theoretical basis for understanding and simulating the role of titin in striated muscle mechanics.

**Key points:**

1. Passive stress relaxation of cardiac muscle follows a power-law decay, a phenomenon that is explained using a theoretical model of dynamic unfolding of globular domains along polymer chain.
2. The theoretical model simulates the behavior of titin, a giant sarcomere protein linking myosin thick filaments to the Z disk and providing passive restoring force during muscle stretch.
3. The theoretical model is able to account the observed effects of calcium on the effective viscoelastic passive mechanics of cardiac muscle.
4. This model provides a theoretical basis for understanding passive visocelastic properties and titin’s role in striated muscle mechanics.

## Introduction

Titin is a large filamentous protein that connects the Z-disk to the myosin thick filament in sarcomeres and functions as a molecular spring, contributing to the passive mechanical properties of striated muscle, as illustrated in Figure 1. Titin also plays a key role in organizing the sarcomeric structure, ensuring the alignment of contractile proteins, and aiding in the transmission of force during muscle contraction. Single-molecule atomic force microscopy reveals that when the titin molecule is stretched, serial globular immunoglobulin-like (Ig) and N2B domains in the titin chain unfold sequentially, resulting in a non-monotonic force-extension relationship [22]. The reversible domain folding behavior is thought to contribute to governing the viscoelastic and hysteresis behaviors in passive stress-strain relationships in muscle [19].

**Figure 1:**
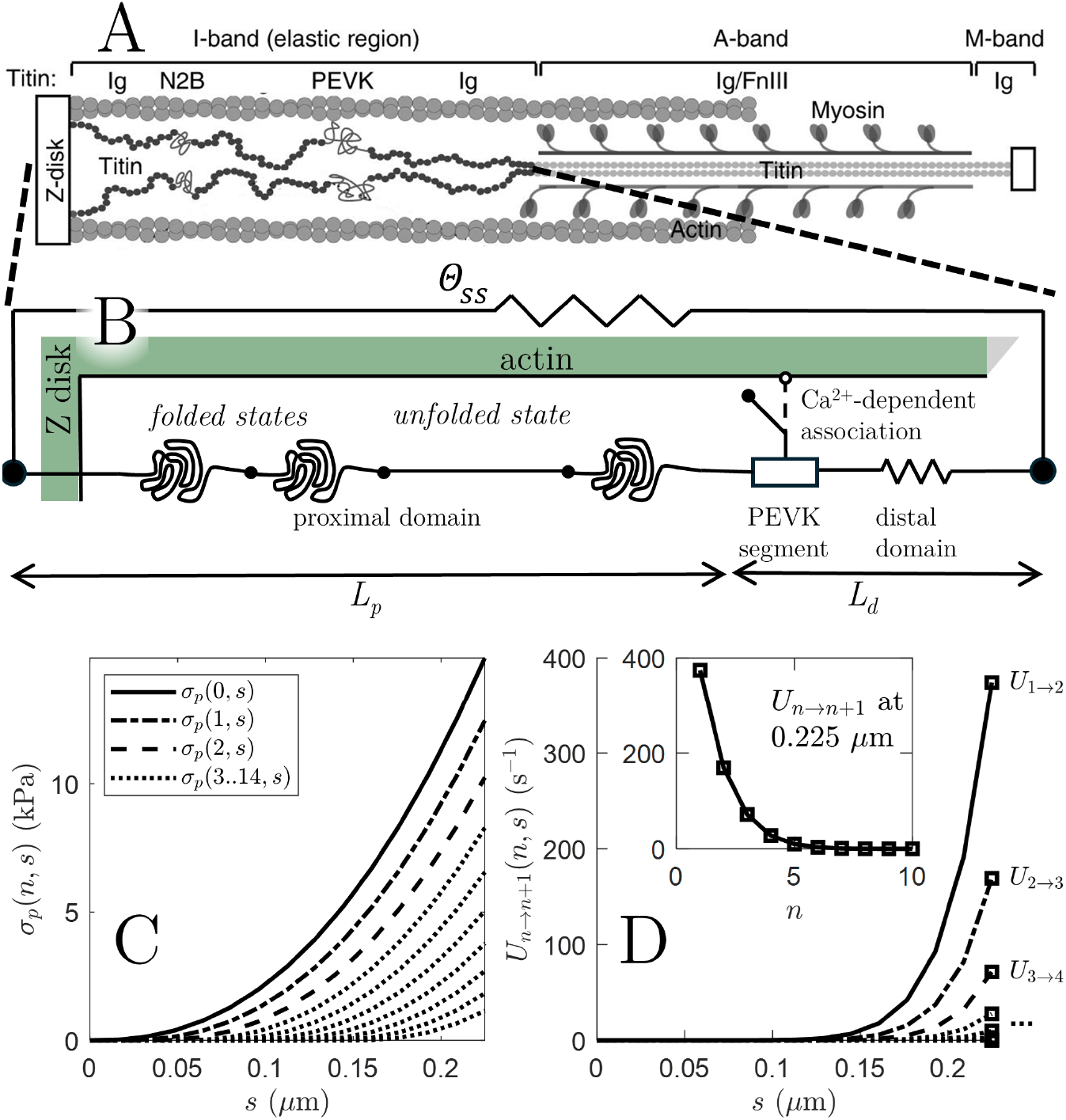
Illustration of titin composition in sarcomere and schematic of model. A. The incorporation of titin into the sarcomere is illustrated, with the titin elastic region in the I-band connecting the myosin thick filament to the Z-disk (adapted from Giganti et al. [5] under CC by 4 international license https://creativecommons.org/licenses/by/4.0/). B. The diagram illustrates the model composition for the titin elastic region. The proximal domain (between the Z-disk attachment and the PEVK domain) is modeled as a series of components (Ig and N2B domains) that can exist in folded or unfolded states. Distal to the PEVK segment is a nonlinear spring labeled “distal domain”. C. The relationship between stress in the proximal domain *σ*_*p*_(*n, s*) and strain of the proximal domain, *s*, is plotted for different values of *n*, the number of unfolded states in a chain. D. The relationship between unfolding rate *U*_*n→n*+1_ and strain of the proximal domain, *s*, is plotted for different values of *n*. Both stress and unfolding rate increase with *s*, and decrease with increasing *n*.

Because titin is the major sarcomeric structural element that provides a restoring force to oppose passive elongation, its mechanical properties are a fundamental determinant of myocardial diastolic function. At sarcomere lengths of roughly 2.1 *μ*m and less, titin is estimated to contribute the majority of the total measured passive tension [7, 10, 26]. Thus the passive pressure-volume relationship associated with diastolic filling of the chambers of the heart is governed, at the cellular and molecular levels, in part by titin mechanics [7, 6, 3]. Phosphorylations of titin residues via PKA, PKD, and PKG kinases in the myocardium are associated with reductions in titin-mediated passive stiffness, while phosphorylation of PKC targets are associated with increased passive stiffness [17, 12, 18]. Thus, the passive mechanical behavior of the myocardium is physiologically regulated via various signalling pathways that regulate the molecular mechanical properties of titin.

Calcium ion concentration is also known to play an important role in regulating the mechanical properties and function of titin. Labeit et al. [16] found the stiffness of the PEVK segment of human titin is sensitive to Ca^2+^, which alters the apparent persistence length associated with the unstressed molecule. Increasing calcium concentration lowers the persistence length or, equivalently, increases the passive elastic resistance to elongation. Kulke et al. [15] showed that the PEVK domain binds to actin in a Ca^2+^-dependent manner. Squarci et al. [24] observed a muscle activation-dependent mechanical “rectifier” behavior, interpreted as the PEVK segment reversibly attaching to the actin fiber in the intact muscle, as illustrated in Figure 1.

Titin’s mechanical behavior arises from unique structural features and functional mechanisms. Its structure is characterized by repeating immunoglobulin-like (Ig) and fibronectin (Fn) domains, and an N2B domain, akin to molecular beads on a string. This modular arrangement allows for extensibility, as certain domains can independently unfold and refold, acting as molecular springs [14, 10, 6]. The extensibility of titin is particularly prominent in the I-band region, which is rich in these domains. Given the fundamental role of titin in determining the passive mechanical properties of muscle, and given the fact that titin mechanical function is dynamically regulated by posttranslational modifications and intracellular calcium, it is not surprising that titin mutations are linked to multiple myopathies [4], including restrictive cardiomyopathy [21], dilated cardiomyopathy[11], and muscular dystrophies [13]. Understanding the molecular-mechanical function of titin is crucial to understanding the mechanisms underlying these mutations.

Baker et al. [1] assessed the Ca^2+^-dependence of the stress response to stretch of permeabilized murine right ventricular trabeculae, showing that addition of Ca^2+^ increases the apparent viscous component of the stress response. To characterize the passive myocardial properties, permeabilized mouse trabecular fibers were stretched over a half-sarcomere lengths from 0.95 to 1.175 *μ*m at ramp speeds ranging from 0.00225 to 2.25 *μ*m·sec^*−*1^, and then allowed to relax for several seconds to minutes. While the rate of tension increase during the stretch phase of the experiment was, unsurprisingly, determined by the speed of the stretch, the rate of tension decay following stretch was, remarkably, observed to following a time-scale independent power-law timecourse. In the presence of calcium the peak tension increased with increasing calcium, even with cross-bridge contraction fully inhibited by para-nitroblebbistatin and mavacamten.

Here, data from Baker et al. are analyzed to develop and identify a theoretical model of the viscoelastic passive mechanical response to stretch in these fibers. The model makes the simplifying assumption of representing passive axial force in the muscle as the sum of two components: a nonlinear elastic component and a visco-elastic component modeled based on the biophysical properties of titin (Figure 1). The model integrates unfolding of a series of globular elements and a calcium-dependent PEVK-actin binding mechanism. The model is able to effectively capture the calcium- and stretch-dependent peak tension as well as the tension decay kinetics observed at different calcium levels. The power-law nature of tension decay is predicted to emerge as a property of the sequence of foldable elements (e.g. Ig and N2B domains), each capable of unfolding and introducing a finite amount of slack into the chain. Equivalently, each unfolding event increases the persistence length of the chain. With each unfolding event, the tension sensed by folded elements in the chain decreases, reducing the effective rate of unfolding. As a result, the time scale of decay increases with decreasing tension during the decay process. The model developed to simulate these processes is a novel tool for investigating passive myocardial mechanics and representing titin mechanics at a mesoscopic scale. It also serves as a means to test and refine hypotheses regarding the function and regulation of passive myocardial properties, and acts as a foundation for integrated modeling of myocardial cell and tissue behavior.

## Methods

### Experimental data on Passive Muscle Mechanics

Data analyzed in this paper are reported in Baker et al. [1]. In brief, right ventricular demembranated trabeculae of 12 week old male (n=3) and female (n = 3) mice were subjected to ramp length extensions from 0.95 to 1.175 muscle length (*L*_0_), where *L*_0_ was established by adjusting muscle length (ML) to set the sarcomere length to 2.0 *μ*m. Muscle dimensions, and active and passive forces were measured at *L*_0_ and at 21°C. The ramp length extension times range from *t*_*r*_ = 0.1 s to 100 s, corresponding to velocites of 2.25 to 0.00225 *L*_0_/s. Following extension, the muscle was held fixed at ML = 1.175*L*_0_ for at least 60 seconds. Muscle stress response to this strain protocol was observed in Ca^2+^-free relaxing solution (denoted pCa= 11), at saturating calcium conditions (pCa= 4.51) to measure maximal stress, and in solution with 50*μ*M para-nitro-blebbistatin and 50*μ*M mavacamten (PNB+Mava), deactivating the cross-bridge force generation. Multiple checks were applied to ensure zero crossbridge attachment. The strain protocol was repeated with PNB+Mava present at five different calcium concentrations: pCa = 4.51, 5.5, 5.75, 6 and 6.25.

An example time course of stress measured of a muscle in relaxed solution (pCa = 11) during this protocol is illustrated in Figure 2, showing that the peak stress is highest for the most rapid ramp speed. Peak stress is approximately 14 kPa for the fastest ramp, and approximately 7 kPa for the slowest ramp. Regardless of peak stress, the stress relaxes to approximately 6 kPa at 60 seconds following the ramp end.

**Figure 2:**
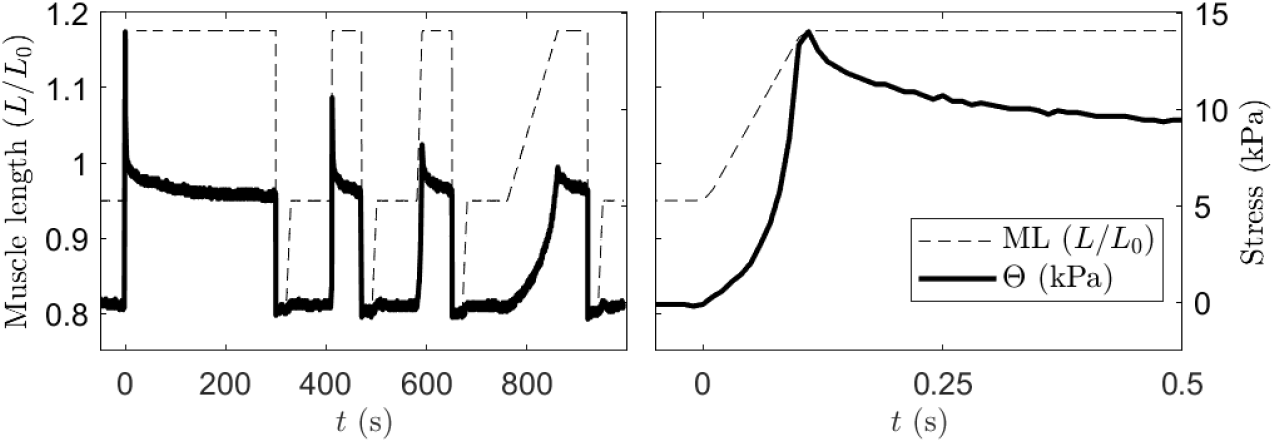
Representative set of measured muscle stress during ramp-up and decay (dataset 1, male). During the ramp increases in length, muscle length (ML) was extended from 0.95*L*_0_ to 1.175*L*_0_ for ramp times of 0.1, 1, 10, and 100 seconds (velocity 2.25×10^*−*3^ - 2.25*L*_0_/s). The right panel shows a detail of muscle length and stress Θ as functions of time for the fastest ramp (0.1 second). Stress increases in a concave-up manner during the ramp increase in length, achieves a peak value at the end of the ramp increase, and then decays.

Measured data were corrected for drift of the force transducer signal by subtracting a spline interpolation of zero-force level, which was assessed by short slack after each ramp. Subsequently, to mimic a normal single cell response, all forces were normalized to maximal force measured in activating solution (maximal [Ca^2+^]) prior to PNB treatment, averaged across multiple experiments and finally rescaled to mean force. This way we avoid error in estimating cross-sectional area and variable fraction of non-contracting actors, as we focus only on passive properties of contractile elements.

### Computational Model

The mathematical model of I-band mechanics takes the form of an ensemble of titin chains, with a single member of the ensemble illustrated schematically in Figure 1. The structure of the model is similar to that of Heidlauf et al. [8, 9]. The single titin chain comprises of proximal unfolding domain (e.g. Fn and Ig-like domains), distal domain and a proline-, glutamic acid-, valine, and lysine-enriched domain, or PEVK segment, connected in series. The total length of the protein is denoted *L* = *L*_*p*_ + *L*_*d*_, with *L*_*p*_ and *L*_*d*_ denoting length in the proximal and distal segments. Strain in the proximal domain denoted *s*, which is defined in reference to a reference muscle length *L*_0_:

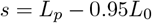

where *L*_0_ = 1 *μ*m is a reference half-sarcomere length, and 0.95*L*_0_ is the initial half-sarcomere length in the ramp extension experiments used to identify the model. The model explicitly tracks the ensemble distribution of strain *s* and number of unfolded domains *n* in the proximal chain. (As a simplifying assumption unfolding events are treated only in the proximal chain.) The titin strand is assumed to attach to and detach from a neighboring actin filament at the PEVK site, located between the proximal and distal chain domains. The actin-bound configuration is referred to as the attached state. The unbound configuration i s referred to as the unattached state.

The variables in the model are listed and summarized in Table 1. The total stress of the muscle arises from contributions from unattached states, attached states, and contributions from parallel non-titin components. The function *p*_*u*_(*n, s, t*) is the ensemble probability density of *n*, number of unfolded states per chain, and *s*, stretch, in unattached chains, such that

**Table 1.**
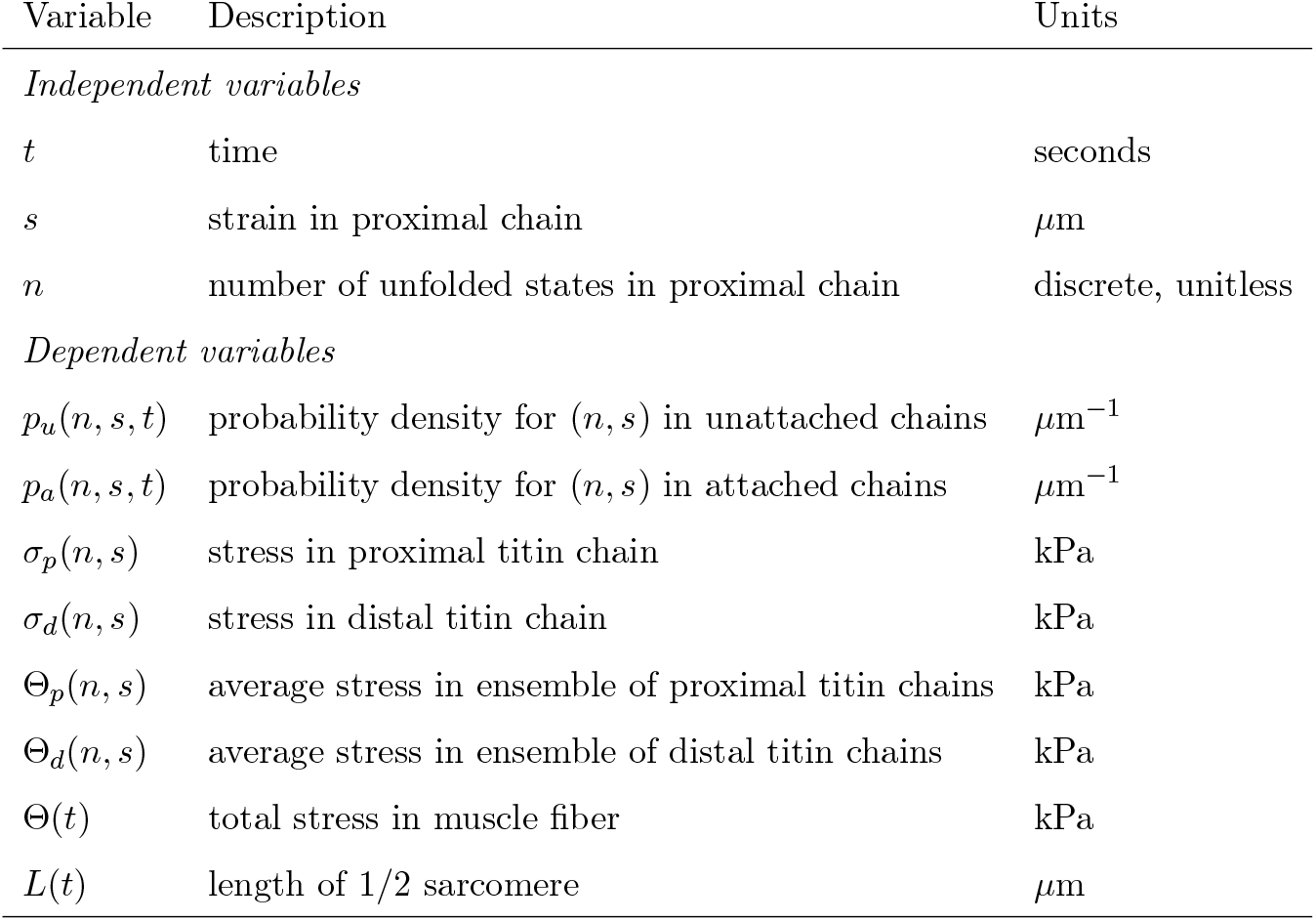
Variables in Model.

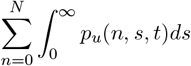

represents the fraction of titin chains that are in the unattached state. Similarly the function *p*_*a*_(*n, s, t*) is the ensemble probability density of *n*, number of unfolded states per chain, and *s*, stretch, in attached chains, such that

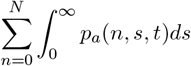

represents the fraction of chains in the attached state. These fractions obey

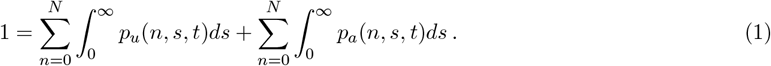

The stress in an individual proximal titin chain domain is assumed proportional to stretch *s*

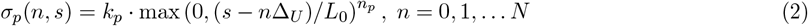

where Δ_*U*_ is the slack length associated with the unfolding of one proximal domain unfolding element, *L*_0_ is a reference half-sarcomere length taken to be 1.0 *μ*m, and *N*_*g*_ is the total number of unfolding domains assumed in the chain. The relationship between *σ*_*p*_(*n, s*) and *s* is illustrated in Figure 1C. The average stress associated with the ensemble of unattached titin chains is computed

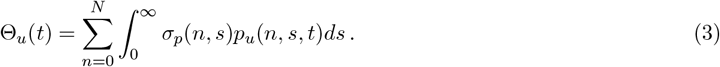

The distal titin domain is assumed to behave like a nonlinear spring. The stress in this element is assumed proportional to stretch of the element *L* − *s* − 0.95*L*_0_:

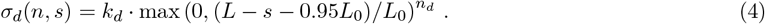

The average stress associated with the ensemble of distal domains is computed

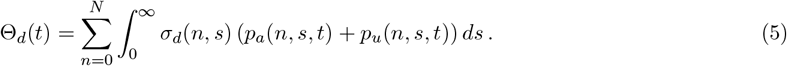

Here titin molecules in both the unattached and attached states contribute to the total force.

Proximal and distal forces in an individual chain are assumed to be balanced, *σ*_*d*_(*n, s*) = *σ*_*p*_(*n, s*). This force balance is achieved in the model by deforming the proximal chain with velocity

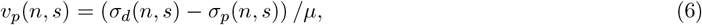

where *μ* is a viscosity factor, that also helps to achieve an effective force balance throughout the simulations.

Unfolding of a single domain, associated with a chain transition from state *n* to *n* + 1, occurs at a rate that is proportional to the strain on the globular chain:

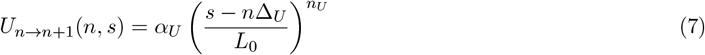

where Δ_*U*_ is the slack associated with one globule unfolding. The unfolding rate *U*_*n→n*+1_(*n, s*) is plotted as a function of *s* in Figure 1D. Refolding events—transitions from state *n* to *n* − 1—are not considered in the current model.

With the rates of stretching and unfolding defined by Equations (6) and (7), the governing equation for *p*_*u*_(*n, s*)—the probability density of *n* and *s* for unattached chain states—is

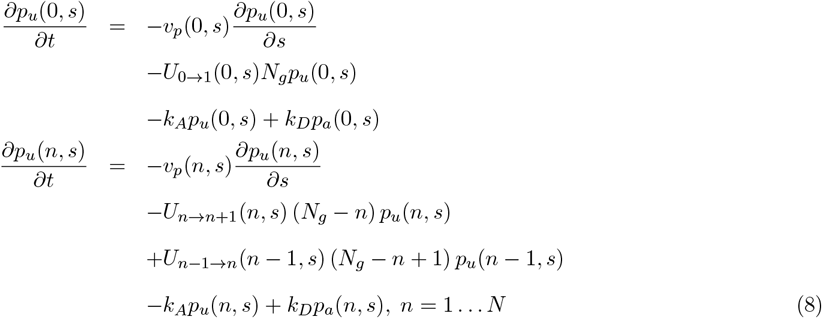

Similarly, the governing equation for *p*_*a*_(*n, s*)—the probability density of *n* and *s* for attached chain states—is

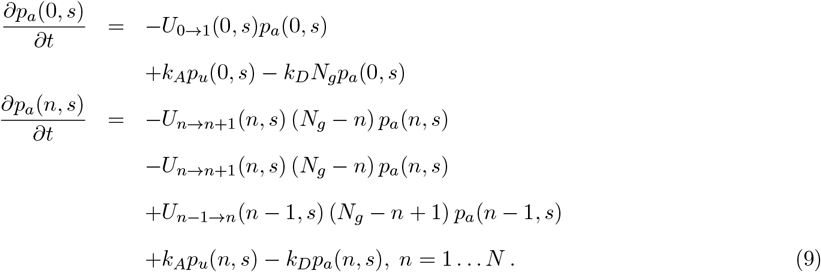

Equations (8) and (9) assume that attachment and detachment of the PEVK domain of the titin chain to actin occur at rates *k*_*A*_ and *k*_*D*_. For calcium-free conditions (pCa = 11) *k*_*A*_ = 0 and no attachment occurs.

Model parameter definitions and estimated values are listed in Table 2. The force proportionality constants *k*_*p*_, the PEVK attachment rate *k*_*A*_ and unfolding rate are assumed to be calcium-dependent. The value of *N*_*g*_ is set to 10, a value that represents the numerical discretization of the chain and not meant to represent the true number of unfolding domains in the proximal chain.

**Table 2.** Adjustable parameters in model.

| Parameter | Description | Value | Units |
| --- | --- | --- | --- |
| *Adjustable parameters (Relaxed, calcium-free, $pCa \approx 11$)* |  |  |  |
| $k_p$ | proportionality constant for proximal chain length-stress relation | 512.3 | kPa |
| $k_d$ | proportionality constant for distal chain length-stress relation | $40 \times 10^3$ | kPa |
| $\alpha_U$ | proportionality constant for unfolding rate | $2.668 \times 10^7$ | $\text{sec}^{-1}$ |
| $n_p$ | exponent for proximal chain length-stress relation | 2.37 | unitless |
| $n_d$ | exponent for distal chain length-stress relation | 2.74 | unitless |
| $n_U$ | exponent for unfolding rate | 9.035 | unitless |
| $\Delta_U$ | slack length associated with unfolding one unit | 0.165 | $\mu\text{m}$ |
| $k_{ss}$ | proportionality constant of the parallel spring | $1.0168 \times 10^9$ | kPa |
| $n_{ss}$ | exponent for the parallel spring | 12.8 | unitless |
| $\mu$ | viscosity of dashpot element | 0.678 | $\text{kPa} \cdot \text{sec} \cdot \mu\text{m}^{-1}$ |
| *fixed parameters* |  |  |  |
| $N_g$ | number of discretized unfolding units in a chain | 10 | unitless |
| $L_0$ | reference half-sarcomere length | 1.0 | $\mu\text{m}$ |
| *Calcium-sensitive parameters (Activated, $pCa = 4.51$)* |  |  |  |
| $\alpha_U$ | proportionality constant for unfolding rate | $2.668 \times 10^7$ | $\text{sec}^{-1}$ |
| $n_U$ | exponent for unfolding rate | 9.035 | unitless |
| $k_A$ | rate of attachment of PEVK element to actin | $5.381 \times 10^{-3}$ | $\text{sec}^{-1}$ |
| $k_D$ | rate of detachment of PEVK element from actin | 0.383 | $\text{sec}^{-1}$ |

The total passive muscle stress is computed from a sum of contributions from titin and from a parallel nonlinear spring that represents the steady state stress at a given length:

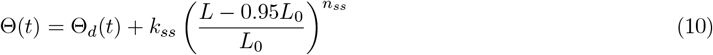

For the ramp extension experiments simulated here the initial state of the model is *p*_*u*_(*n, s*) = *δ*(*s*)*δ*_0,*n*_ and *p*_*a*_(*n, s*) = 0, assuming no attachment to actin with zero strain and all globular elements in the folded state. Prior to initiating the ramp-up, the model is run to reach an initial steady state. Computer codes for simulating the model are available at https://github.com/beards-lab/TitinViscoelasticity.

### 2.1 Parameter identification

A global optimization was performed to find the best match to the measured tension responses to ramps across all the different calcium levels. The objective of this optimization process was to simultaneously minimize the least squares differences between model outputs and data and the number of parameters that are assumed to vary with calcium concentration. In practice we first identified the model for the high-calcium case and then a depth-first recursive search was performed to explore all possible parameter subsets that could be adjusted to match the low-calcium data. Out of all possible subsets, we found that at least four parameters are needed for a satisfactory fit. Although there were five plausible combinations of four parameters, each subset contained proximal segment stiffness *k*_*p*_, one of PEVK attachment or detachment rates (*k*_*A*_ or *k*_*D*_) and either a term affecting unfolding rate (either *α*_*U*_ or *n*_*U*_) or proximal chain length-stress exponent *n*_*p*_. The lowest-cost combination of four parameters was *k*_*A*_, *k*_*p*_, *α*_*U*_ and *n*_*U*_, which were varied as functions of calcium concentration. To fit the intermediate calcium levels, we adjusted these four parameters only, while enforcing monotonicity with calcium concentration.

## Results

### Properties of the stress relaxation

Average stress time-course data from experiments of Baker et al. [1] at pCa = 11 are plotted in Figure 3 on linear, semilog-y, and log-log scales. To plot the data on semilog-y and log-log scales a common long-time steady-state stress level Θ_∞_ was estimated and the data were shifted in time so that the decaying tails of the stress data are optimally overlaying. For these data the estimated value of Θ_∞_ is 4.84 kPa, and the total stress minus the steady-state stress, Θ − Θ_∞_ is plotted on the y-axes in the semilog and log-log panels in Figure 3. The time shifts used to overlay the decay tails are indicated in the figure.

**Figure 3:**
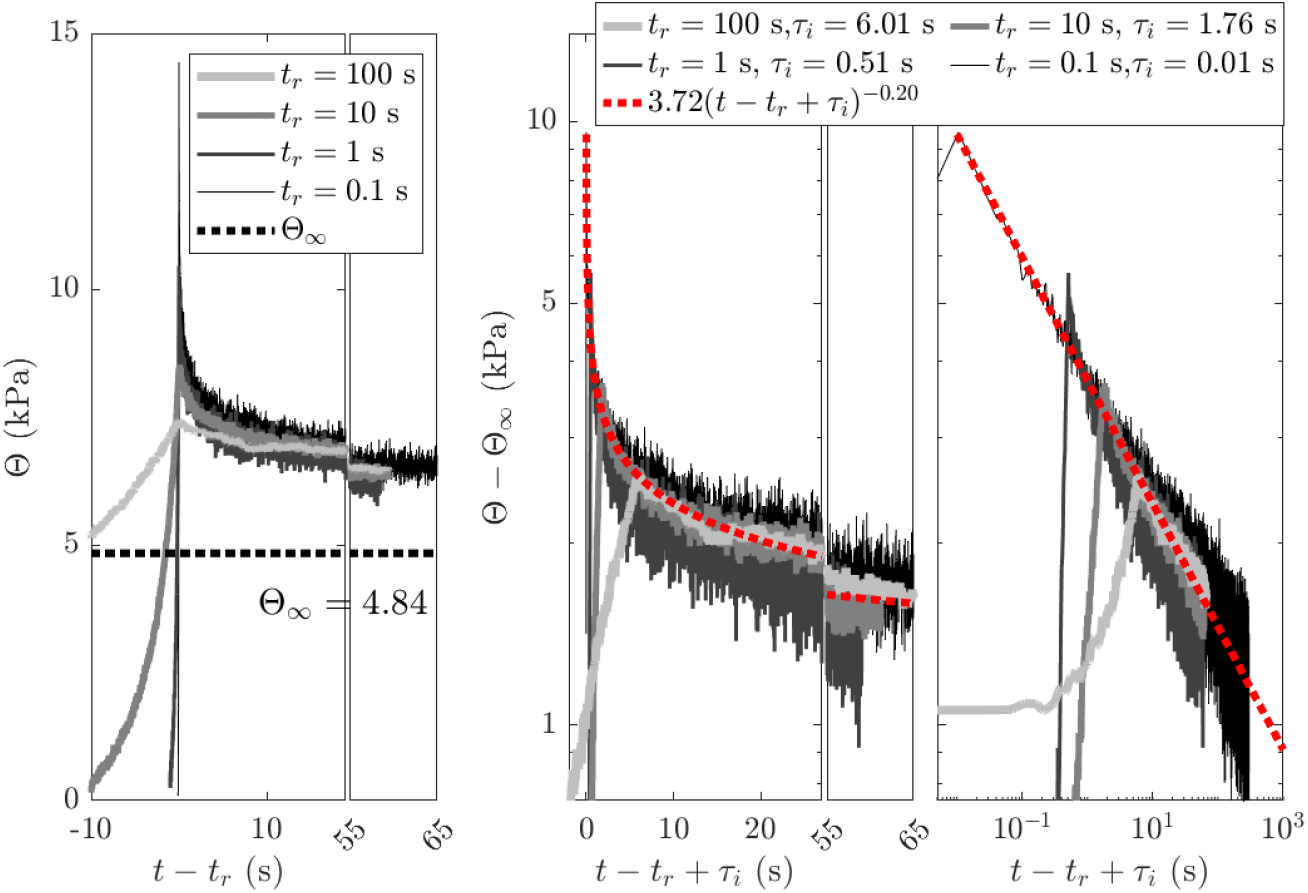
Analysis of measured stress decay in relaxed muscle fibers. The left panel plots stress versus time for the four ramp speeds on a linear scale. The middle and right panels plot stress versus time for the four ramp speeds on semilog-y and log-log scales. The semilog and log-log plots show a power-law function fitting of the decay curves for all ramps. The ramp times *t*_*r*_ = 0.1 s to 100 s and time shifts *τ*_*i*_ for the semilog and log-log plots are indicated in the figure inset.

Figure 3 illustrates that the stress decay process does not follow exponential kinetics (does not follow a straight line on the semilog-y plot). Rather, the decay follows a power law with Θ ∼ *t*^*−*0.20^. This power-law decay is independent of the ramp speed or peak height. Moreover, the observed power-law decay, with an exponent of −0.20, suggests that stress decay may extend over extraordinarily long time scales. This result also compares remarkably well with behavior observed for human myocardium, which shows a power-law decay, with an exponent of −0.184 [20]. As illustrated in the figure, the extrapolation of the power-law decay predicts apparent viscous contribution of 1 kPa to the passive stress 1000 seconds after the peak stress, while a decrease to 0.1 kPa at this rate would theoretically take over 800 years. Thus, the estimated Θ_∞_ of 4.84 kPa is lower than the apparent steady-state stress observed 60 seconds after the peak stress reported by Baker et al. [1]. In contrast, calcium-treated muscle shows a more complex response that cannot be fit using power-law behavior alone (see below).

### Model analysis of passive muscle dynamics under Ca^2+^-free conditions

Figure 4 shows optimal model fits to stress data collected under calcium-free (pCa = 11) conditions. The ramp times *t*_*r*_ indicated in the figure denote the length of time each ramp takes. The greatest peak stress is associated with the fastest ramp, *t*_*r*_ = 0.1 seconds. The computational model is able to correctly match not only the peak heights for different ramp durations, but also the stress increase during ramp up and the decay time course. The right panel of Figure 4 shows that the model is also able to capture the observed power-law decay in stress, with stress decay in the model following Θ ∼ *t*^*−*0.24^ over three decades in time.

**Figure 4:**
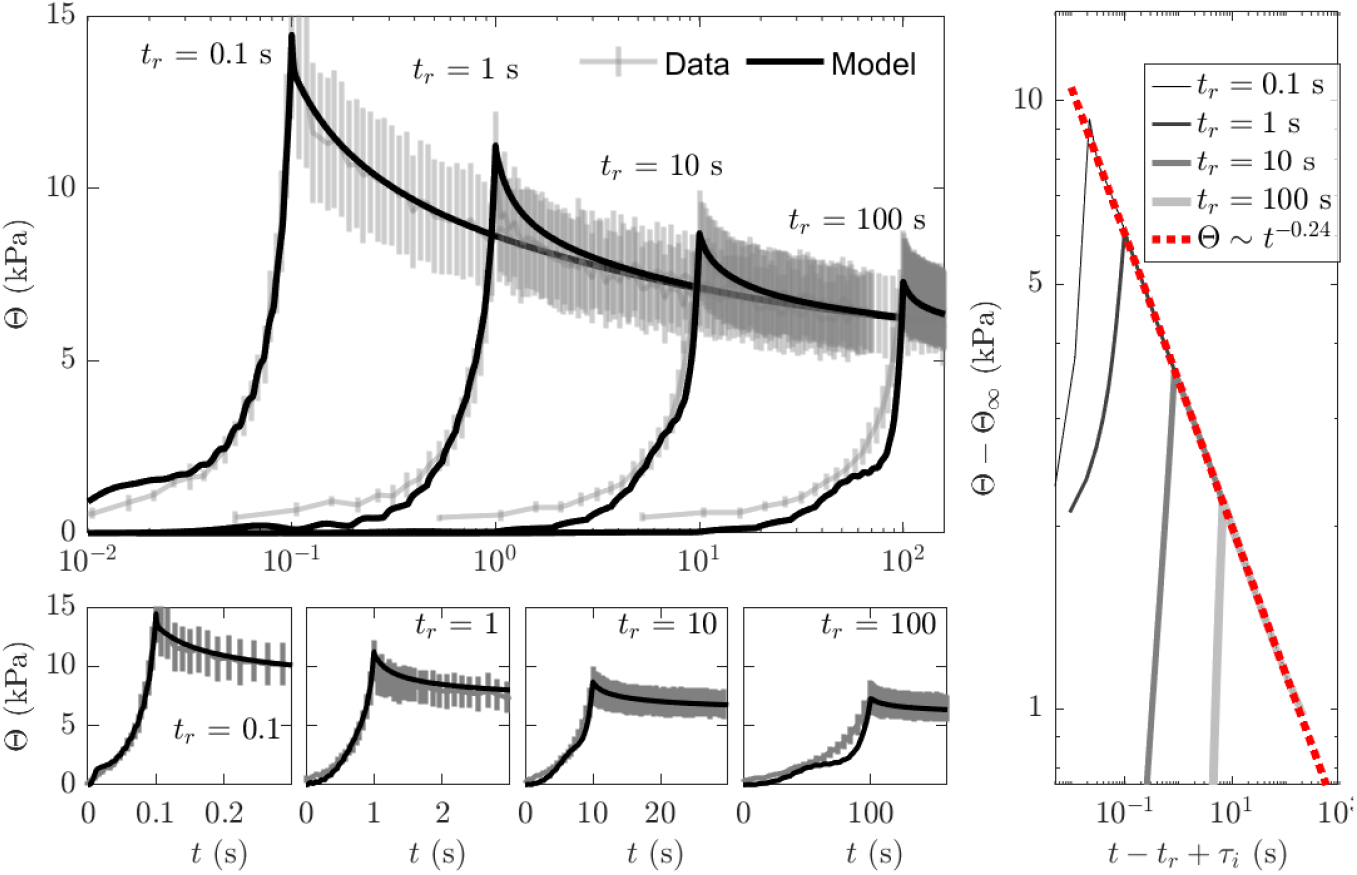
Model fit to stress data following ramp increases in muscle length at zero calcium (pCa = 11). A. For all time courses the muscle is lengthened from 0.95*L*_0_ to 1.175*L*_0_ in a linear ramp and then held at the final length to observe the stress decay. The top panel show stress data and optimal model fit on a semilog-x plot. The bottom panel shows the peak and initial stress decay on linear scale plots for each ramp time, from *t*_*r*_ = 0.1 seconds to *t*_*r*_ = 100 seconds. B. Model-predicted stress decay is shown on a log-log plot illustrating the power-law behavior Θ ∼ *t*^*−*0.24^.

The probability state space of the simulated ensemble of elastic chains, *p*_*u*_(*n, s*), during the 1-second ramp stretch and decay is illustrated in Figure 5, which shows the state probability as heat maps in the (*n, s*) space at different time points in the experiment. In the initial state all chains are folded, there is no strain in the system, and thus *p*_*u*_(*n, s*) = *δ*(*s*)*δ*_0,*n*_. When chain is rapidly stretched by 0.225 *μ*m, at time *t* = 0.1 seconds, *p*_*u*_(*n, s*) has a peak at approximately *s* = 0.18 *μ*m and *n* = 2, meaning that the proximal part of the chain is stretched by approximately 0.19 *μ*m in average and the distal part by 0.03 *μ*m. At this time point there is also nonzero probability at *n* = 1 … 4, meaning that some of the chains in the simulated ensemble have already begun to unfold. During the relaxation phase of the experiment the peak in the probability density shifts to higher values of *n*. At the final time point in the simulation (*t* = 100 seconds), the peak of *p*_*u*_(*n, s*) is at *n* = 9 and *s* ≈ 0.22 *μ*m, associated with a strain in the proximal chain of 0.22 *μ*m with 9 globular units in the chain unfolded. Since each unfolding introduces Δ _*U*_ slack in the chain, the final state is associated with roughly 9Δ_*U*_ ≐ 0.15 *μ*m of slack in the chain.

**Figure 5:**
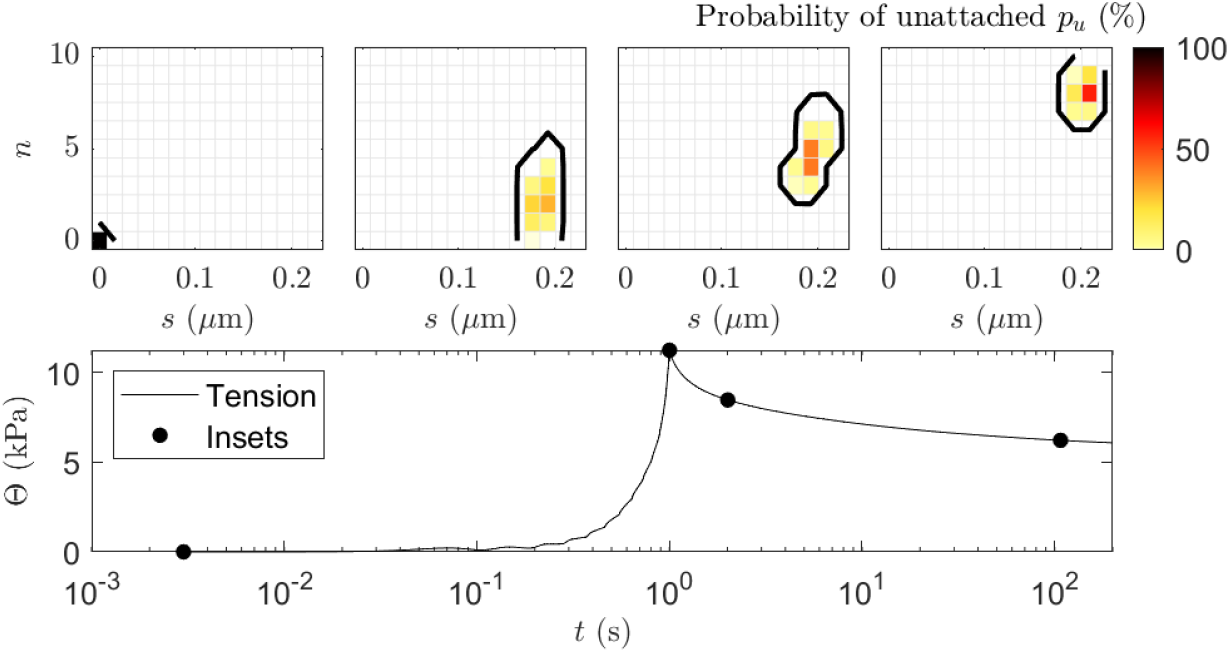
Probability space of simulated chain at no calcium. Top: the probability density *p*_*u*_(*n, s*) is illustrated as a heat map, with contours drawn at 1%, at four different times in the simulation of the fast ramp extension experiment for the low-calcium condition. The bottom panel shows the simulated stress time course and the four time points (*t* = 10^*−*3^, 1, 2, and 100 seconds) for which the the probability density *p*_*u*_(*n, s*) is illustrated in top panels as a per cent of all states. There is no attachment considered at minor calcium concentrations, thus *p*_*a*_(*n, s*) is always zero and is not shown.

### Model analysis of passive muscle dynamics under high-Ca^2+^ conditions

Figure 6 shows optimal model fits to stress data collected under high-calcium (pCa = 4.51) conditions. Similar to the stress response at low calcium, the greatest peak stress is associated with the fastest ramp, *t*_*r*_ = 0.1 seconds. The peak stresses at each extension speed are all markedly higher at pCa = 4.51 compared to peak stresses at pCa = 11. The model captures this Ca^2+^-induced increase in peak force through the Ca^2+^-dependence of chain stiffness, unfolding rate and PEVK domain attachment to actin. The simulated stress decay time courses in a log-log scale in the right panel of Figure 6 show that predicted long-time decay behavior at high calcium follows the same power-law behavior at low calcium.

**Figure 6:**
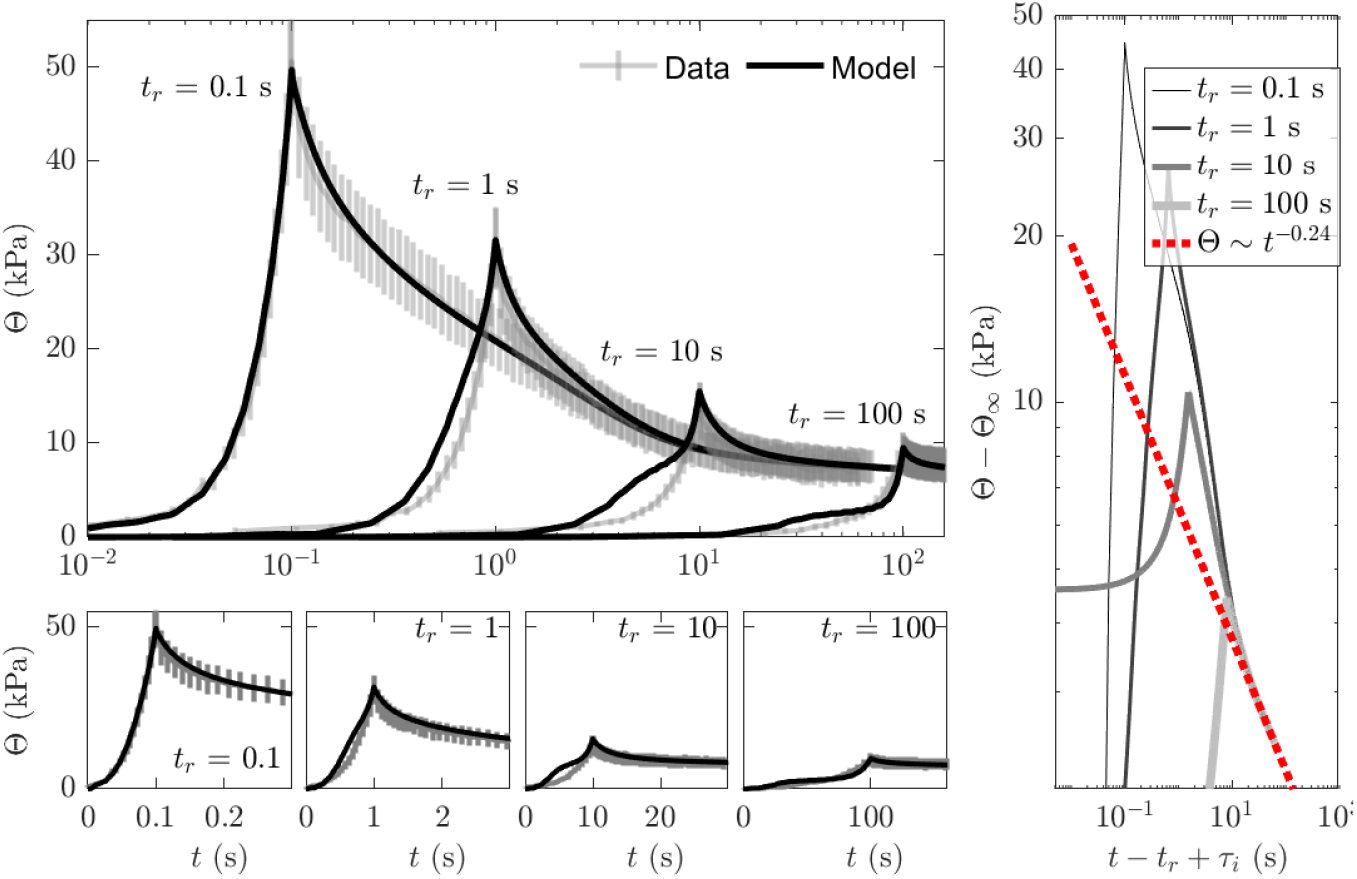
Model fit to stress data following ramp increases in muscle length at high calcium (pCa = 4.51). For all time courses the muscle is lengthened from 0.95*L*_0_ to 1.175*L*_0_ in linear ramp and then held at the final length to observe the stress decay. The top-left panel show stress data and optimal model fit on a semilog-x plot. The bottom panels show the peak and initial stress decay on linear scale plots for each ramp time, from *t*_*r*_ = 0.1 seconds to *t*_*r*_ = 100 seconds. Right: Model-predicted stress decay is shown on a log-log plot, with second-half of the decay (30–60s) fitted with a power law.

At high calcium the simulated titin elastic chain adopts conformation in both the *p*_*u*_(*n, s*) and *p*_*a*_(*n, s*) spaces, as illustrated in Figure 7, which shows the state probability as heat maps in the (*n, s*) space at different time points in the experiment. At time *t* = 10^*−*3^ seconds, the binding and unbinding to actin are in an equilibrium balance, with approximately 1.5% of conformations in the attached state. In addition, in the initial state all chains are folded with no strain on the titin chain, and thus *p*_*u*_(*n, s*) + *p*_*a*_(*n, s*) = *δ*(*s*)*δ*_0,*n*_. When chain is rapidly stretched by 0.225 *μ*m, at time *t* = 1 seconds, the peak in *p*_*u*_(*n, s*) moves to *s* ≈ 0.16 *μ*m, predicting that the proximal titin chain is relatively less stretched at this time point in the simulation compared to behavior at low calcium (Figure 5). In contrast to the calcium-free response, the stiffness is larger, but the unfolding rate is faster. During the initial phases of unfolding, the attached states increase the tension by holding on previously attached state (here 3rd left middle panel - around 1% of the states are still attached at zero strain, thus 1% of the distal chain have to have *L*_*d*_ = *L*_*max*_). In later decay the attached states reattach (or unfold) and drift to the same strains as the unattached, reducing the PEVK attachment contribution. Figure 11 (in the Appendix) shows simulations of the high-calcium (pCa = 4.51) with PEVK binding disabled. These simulations predict the PEVK binding is an important factor in the viscoelastic response and that it is the actin-titin interaction at high calcium.

**Figure 7:**
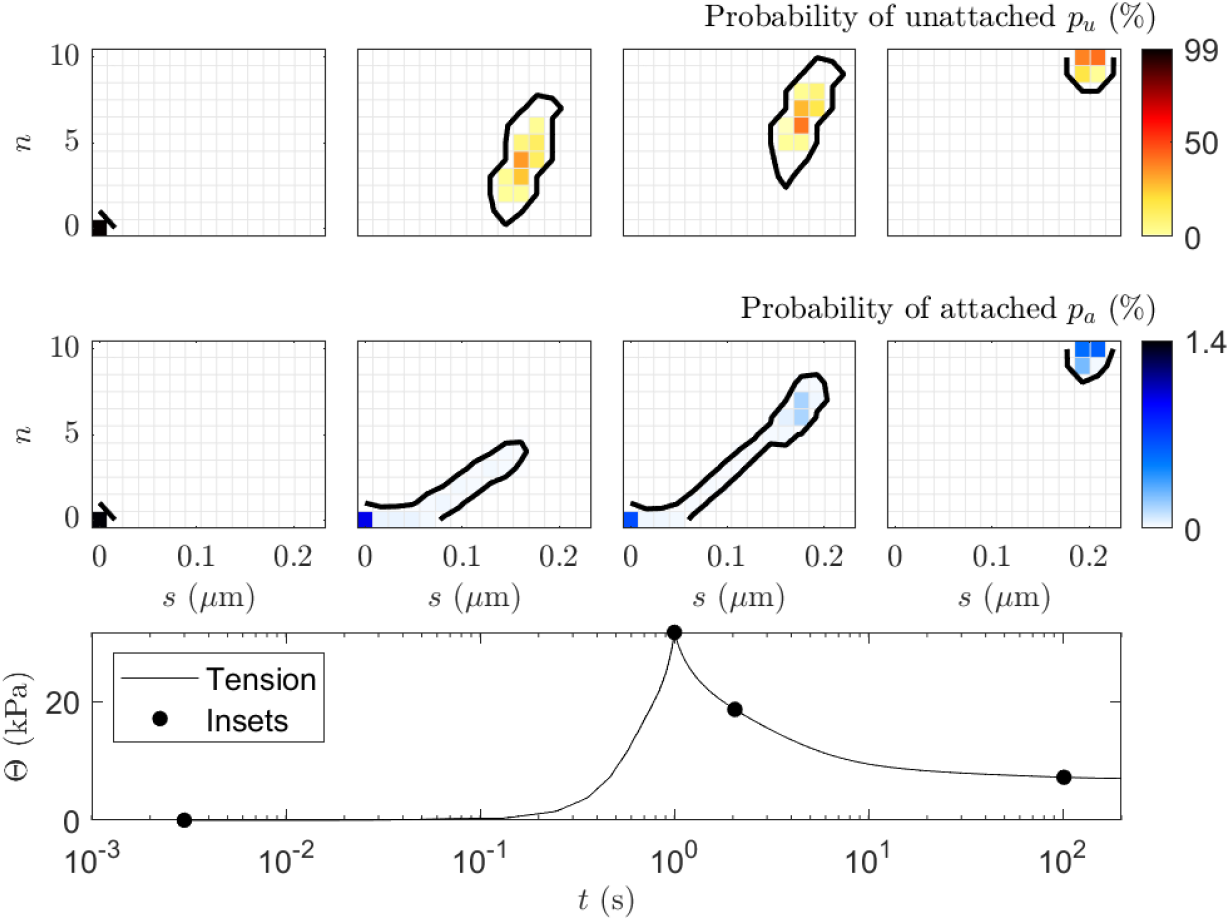
Probability space of simulated chain at high calcium (pCa = 4.51). Top: the probability density of the unattached states *p*_*u*_(*n, s*) is illustrated as a heat map, with contours drawn at 1%, at four different times in the simulation of the 1s ramp extension experiment. Middle: the probability density of the attached states *p*_*a*_(*n, s*) is illustrated as a heat map, at four different times in the simulation of the fast ramp extension experiment. The bottom panel shows the simulated stress time course and the four time points (*t* = 10^*−*3^, 1, 2, and 100 seconds) for which the probability densities *p*_*u*_(*n, s*) and *p*_*a*_(*n, s*) are illustrated in top and middle panels as a per cent of all state probabilities.

Figure 8 shows stress decay after a ramp-up, as measured by Baker et al. [1], and simulated using the proposed model over a range of calcium concentrations, demonstrating that the model effectively captures the calcium-sensitive mechanisms. The peak stress gradually increases from less than 20 kPa at low calcium (pCa = 6 to around 50 kPa at pCa = 5.75, indicating high senstivity to [Ca^2+^] in the micromolar range. With decreasing calcium concentration the stress decay tends to be more similar to a power law, as discussed above. Model parameters across a range of calcium concentrations are well fit using a Hill curve (Figure 9). The estimated Hill coefficients for the calcium dependence of these four parameters are remarkably similar: *K*_*A*_ = 5.87 ± 0.09.

**Figure 8:**
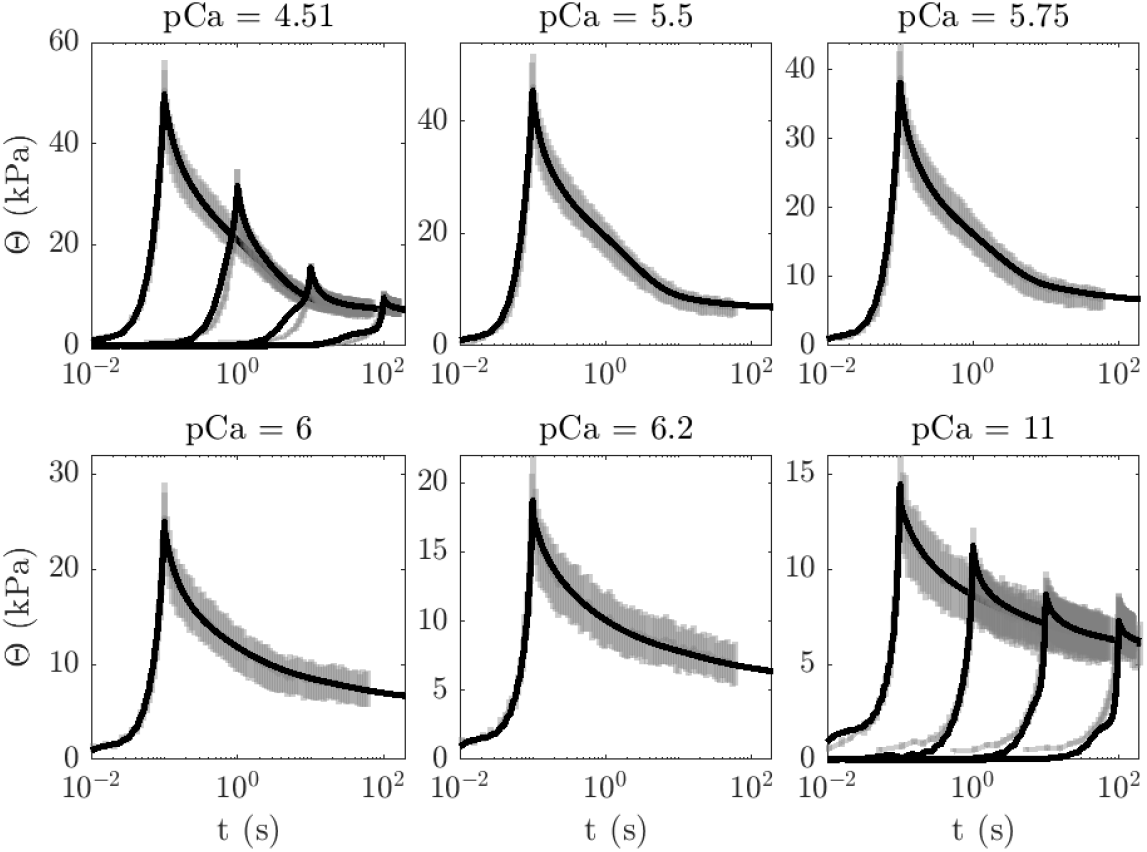
Comparison of measured and simulated stretch response across a range of calcium concentration. For intermediate calcium concentrations only the fastest ramp data were acquired.

**Figure 9:**
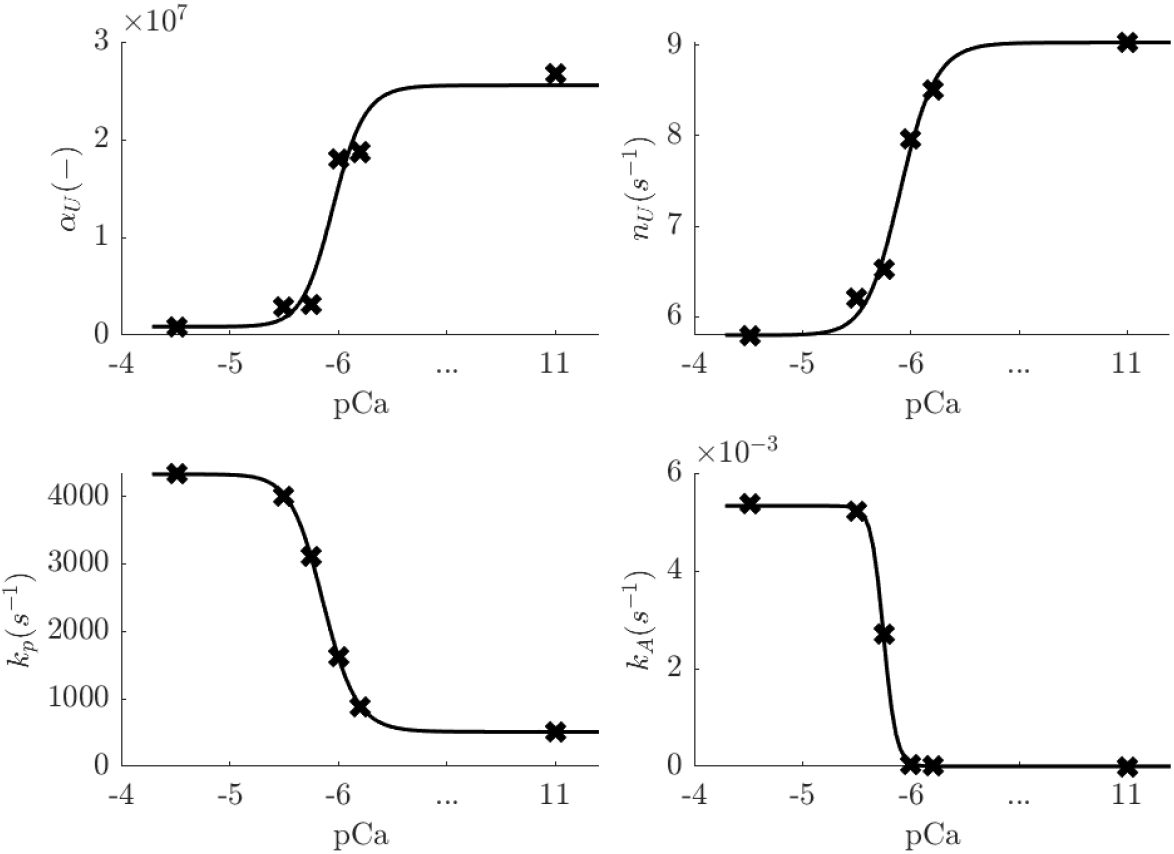
Parameters varied with Ca^2+^ concentration required to fit ramp responses over a range of Ca^2+^ concentrations shown in Figure 8. Parameters were identified allowing only monotonous transition and were subsequently fit using Hill curve.

### Simulation of cyclic loading

To explore preconditioning behavior we simulated stress response to cyclical sinusoidal loading, with the initial condition of the model in the fully folded state: *p*_*u*_(*n, s*) = *δ*(*s*)*δ*_0,*n*_. Cyclical loading was simulated by imposing a half sarcomere length of *L*(*t*) = *L*_0_ + (*A/*2)(cos(*ωt* − *π*) + 1), with amplitude *A* = *L*_*max*_ = 0.225 *μ*m and frequency *ω/*(2*π*) = 1 s^*−*1^. Predicted stress response to this length time course is illustrated in Figure 10 for the zero-calcium condition. Simulations show that during the initial extension stress increases to a peak of approximately 12 kPa. The peak stress decays with each subsequent cycle of stretch. The right panel of the figure illustrates this hysteresis behavior in a plot of stress versus stretch.

**Figure 10:**
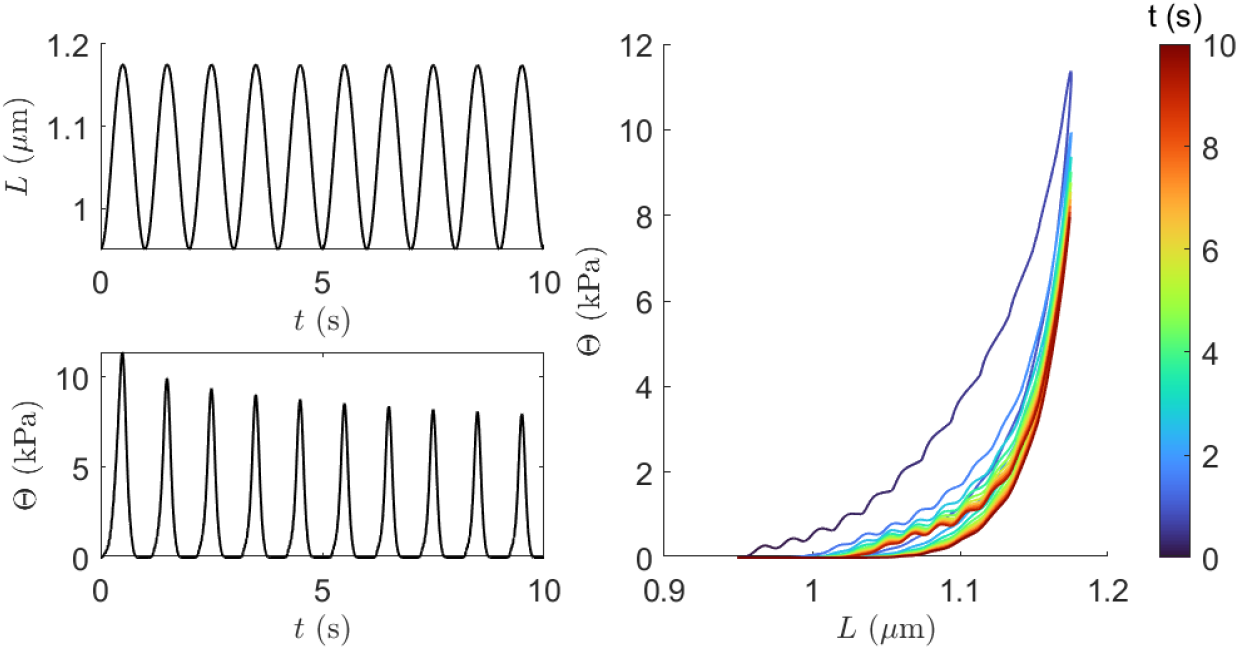
Simulation of cyclical loading at zero calcium. The left panel shows the imposed half-sarcomere length *L*(*t*) and resulting model-predicted stress Θ(*t*) for loading frequency of 1 Hz. The right panel shows the predicted length-stress relationship demonstrating loading-unloading difference and preconditioning of the relaxed muscle. The non-smooth behavior seen in the stress-strain plot is a numerical artifact of the state space discretization.

The predicted stress-strain behavior, showing a drop in peak force from one cycle to the next, is similar to experimental data, for example, from biaxial extension of myocardium [23] or [19]. The ability to capture this behavior suggests that the model provides a mechanistic basis for simulating preconditioning phenomena in striated muscle tissue. Unlike experimental results on intact tissue, the simulated hysteresis loop collapses after several loading cycles. This is because the current model does not account for refolding of unfolded domains in the chain. Future investigations into titin-mediated preconditioning and hysteresis behavior will require the current model to be extended to account for refolding events.

## Discussion

We developed and analyzed a mathematical model of the passive mechanics of myocardium represented with an elastic component and a component representing titin, in which titin chains are represented as an ensemble of chains with a probabilistic distribution of strain and degree of unfolding of serial elements in the chains. The model captures a broad range of important phenomena over multiple calcium concentrations that contribute to the nonlinear passive mechanics of myocardial tissue.

In fitting model output to data from Baker et al. [1] the model accurately reflects the observed stress response to a ramp increase in strain, with a concave-up increase in stress during the extension, followed by a decay. As observed by Baker et al. [1], the peak stress achieved in the ramp extension experiment increases with increasing extension speed, reflecting an apparent viscous component of the stretch response. This phenomenon is predicted to be associated with the dependency of rates of serial titin domain unfolding events on strain. At relatively slower extension rates, unfolding domains have more time to unfold during extension, increasing the persistence length of the chain, and resulting in less stress compared to stress at higher extension rates. Regardless of extension speed, when the muscle is held at a fixed length following a ramp extension, the stress decays with a time course that follows *t*^*−β*^, where *β* is estimated to be 0.2 from the raw experimental data and 0.24 from model simulations. This power-law decay emerges as a property of the sequence of globular elements in the titin chain. Each element is capable of unfolding and introducing a finite amount of slack into the chain. Equivalently, each unfolding event increases the persistence length of the chain. As each unfolding event introduces slack into the chain the stress sensed by unfolded elements in the sequence decreases, reducing the effective rate of unfolding. Thus, the effective rate of decay decreases as the decay proceeds, resulting in the slow power-law decay.

This unfolding of serial foldable elements in the titin chain is also predicted to contribute to the hysteresis and preconditioning in the muscle stress-strain relationship. The mechanism underlying hysteresis and preconditioning phenomena in the model follows from the fact that the effective stiffness in the elastic I band region is greatest in the fully folded state. Muscle stretching leads to unfolding, which reduces the effective stiffness of the muscle.

In sum, the unfolding beads-on-a-chain titin model captures the following features of passive myocardial mechanics:

1. The dependency of maximum stress on the rate of muscle extension—the apparent viscous component of the stress response—in a muscle-stretch experiment, as illustrated in Figures 4, 6, and 8.
2. The power-law kinetics of stress relaxation, as illustrated in Figures 3 and 4.
3. Preconditioning in the passive viscoelastic muscle stress-strain behavior, as illustrated in Figure 10.

Furthermore, by including the calcium-dependent PEVK domain attachment to actin and calcium-dependent chain stiffness, the model captures the observed dependence of peak stress on calcium concentration, as illustrated in Figure 6. This calcium dependence may be important *in vivo* as calcium stimulates cross-bridge activation and force generation. The model is able to match the high-calcium-treated muscle response by stiffening the proximal domain, destabilizing domain unfolding, and attaching PEVK to actin. Although only small fraction of PEVK are predicted to be attached (see Figure 7), this attachment makes an observable difference, especially in the fastest ramp-up (compare to Figure 11 of the Appendix). This interpretation is in line with observations of Squarci et al. [24].

### Model limitations

The model formulation invokes a number of simplifying assumptions, including the simplification that domains may unfold only in the titin segment proximal to the PEVK domain. Moreover the model does not distinguish specific unfolding domains, such as Ig, Fn, and N2B elements, which may be dominant at lower stresses [25]. This validity of this simplification is apparent in the ability of the model to match the observed stress relaxation data. However, to represent different behaviors associated site-specific mutuations, domain deletions, or post-translational modification, a more detailed model formulation may be required. Since the attachment of the PEVK domain to actin does not come into play in low-calcium (pCa = 11) conditions, the assumption that unfolding events occur in only one segment of the chain does not influence the behavior of the model at zero or very low [Ca^2+^] and incorporating distal domain unfolding would make the model more complex.

For applications demonstrated here the model does not account for refolding of unfolding domains in the titin chain. However to more accurately simulate hysteresis phenomena, as well as to simulate the contribution of titin mechanics to passive myocardial mechanics *in vivo* it would be necessary to incorporate refolding into the model. Preliminary tests suggest adding a constant refolding rate results in hysteresis similar to those observed experimentally, but a robust incorporation of refolding kinetics would require additional experiments.

Although model parameter identification is based on a set of relatively rich and informative data sets, the estimated parameter values likely do not represent a unique global solution. This lack of confidence in the global uniqueness of the parameter estimates does not mean that the predictions of the model are flawed, but rather that the parameters are not necessarily well identifiable and their values should be considered valid only in mutual combination and within the context of this model.

The experiments used to identify the model were conducted at 21°C. In future it would be valuable to attain relevant data at 37°C to investigate how the mechanical phenomena explored here depend on temperature.

In the proposed model, the increased tension with increased calcium concentration is attributed to (1.) PEVK attachment to actin, (2.) stiffening of proximal element, and (3.) destabilizing unfolding rate (increasing rate and rate exponent). For other possible candidates (with slightly worse fit, data not shown) it was sufficient to vary (1.) and (2.) and proximal and distal chain exponents (*n*_*p*_ and *n*_*d*_), but keeping the unfolding rate constant. With given data we cannot effectively rule these possibilities out, so we simply used the lowest cost combination of calcium dependencies.

### Attribution of apparent viscous component to titin

In this work we developed a model of viscoelastic passive muscle mechanics based on model of titin as a series of elements that unfold in a stress/strain-dependent manner in parallel with a nonlinear elastic element. Yet analysis of mechanics data obtained under myofilament-extracted conditions from Baker et al. [1] suggests that a component of the apparent viscoelastic response is due to non-myofilament structures. Our analysis of the data from Baker et al. predicts that the parallel elastic component contributes roughly 5 kPa to the peak stress of approximately 15 kPa observed at the fastest ramp speed observed under zero-Ca^2+^ conditions. Of the remaining ≈ 10 kPa *viscous* component, as much as 4 kPa may be attributed to non-myofibrillar structures. Thus, the apparent viscous component of the stretch response at low low Ca^2+^ is not expected to be entirely due to titin. Indeed there exist additional passive mechanical mechanisms that are not included in this model, including an effective viscoelasticity of microtubules [2]. Nevertheless, the general form of our theoretical model—of a linked series of domains that undergo stress-mediated unfolding—may equivalently represent contributions from non-titin mediated processes to the power-law stress relaxation of the myocardium. Moreover, as demonstrated in Baker et al., the Ca^2+^-dependent component of the viscous response is entirely attributable to myofibrillar structures.

## 3 Acknowledgements

This work was supported by Department of Veterans Affairs Merit Review Award I01BX000740 (AJB) and National Heart, Lung and Blood Institute Grants R01 HL154624 (A.J.B., D.A.B.) and R01 HL173346 (D.A.B.)

## 4 Data availability

All data and codes to reproduce figures in this paper are available at https://github.com/beards-lab/TitinViscoelasticity.

## Appendix

Figure 11 shows simulations of the model under high-calcium conditions (pCa = 4.51), but with the PEVK attachment to actin disabled. Simulations predict that even though only approximately 1% of PEVK domains are predicted to be attached, PEVK attachment is associated with a substantial contribution to the stress achieved at the fasted ramp speed at high calcium.

**Figure 11:**
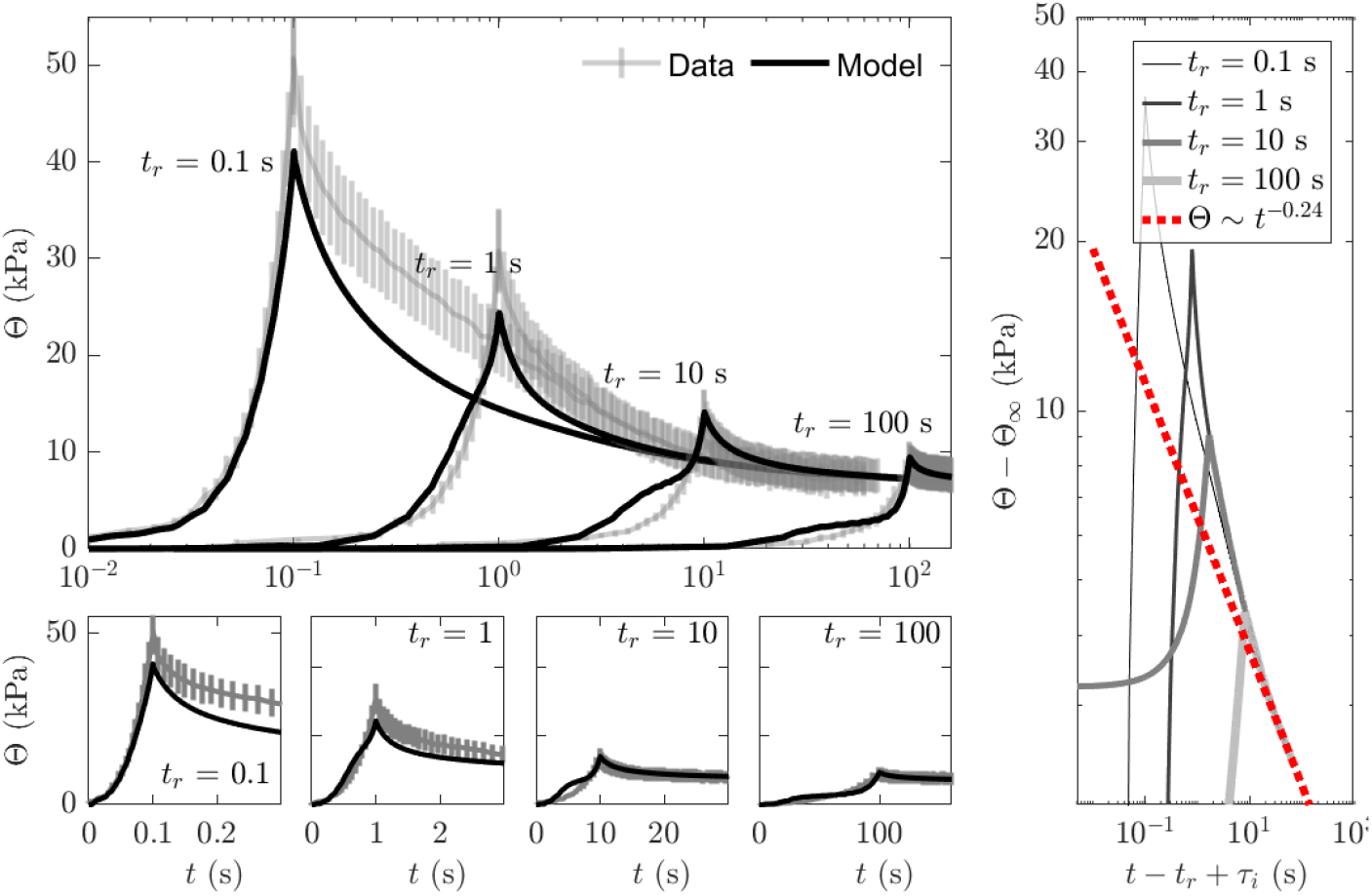
Model simulations of stress response following ramp increases in muscle length at high calcium (pCa = 4.51) with PEVK binding disabled. Simulations are equivalent to simulations shown in Figure 6, but with *k*_*A*_ = 0, resulting in no attachment of the titin to actin.

